# Dr. Sim: Similarity Learning for Transcriptional Phenotypic Drug discovery

**DOI:** 10.1101/2021.09.23.461458

**Authors:** Zhiting Wei, Sheng Zhu, Xiaohan Chen, Chenyu Zhu, Bin Duan, Qi Liu

## Abstract

Transcriptional phenotypic drug discovery has achieved great success, and various compound perturbation-based data resources, such as Connectivity Map (CMap) and Library of Integrated Network-Based Cellular Signatures (LINCS), have been presented. Computational strategies fully mining these resources for phenotypic drug discovery have been proposed, and among them, a fundamental issue is to define the proper similarity between the transcriptional profiles to elucidate the drug mechanism of actions and identify new drug indications. Traditionally, this similarity has been defined in an unsupervised way, and due to the high dimensionality and the existence of high noise in those high-throughput data, it lacks robustness with limited performance. In our study, we present Dr. Sim, which is a general learning-based framework that automatically infers similarity measurement rather than being manually designed and can be used to characterize transcriptional phenotypic profiles for drug discovery with generalized good performance. We evaluated Dr. Sim on comprehensively publicly available *in vitro* and *in vivo* datasets in drug annotation and repositioning using high-throughput transcriptional perturbation data and indicated that Dr. Sim significantly outperforms the existing methods and is proved to be a conceptual improvement by learning transcriptional similarity to facilitate the broad utility of high-throughput transcriptional perturbation data for phenotypic drug discovery. The source code and usage of Dr. Sim is available at https://github.com/bm2-lab/DrSim/.

## Introduction

Genome-wide expression profiles that identify genes showing expression patterns related to a specific phenotypic trait have been successfully applied in drug discovery, such as by elucidating the mechanisms of action (MOAs) for little known compounds or suggesting new indications for existing drugs [1-3]. Based on this concept, various resources, such as the Library of Integrated Network-Based Cellular Signatures (LINCS), have produced massive transcriptional profiles by treating various cancer cell lines with different compounds (also referred to as perturbations) under different conditions [4, 5]. The related processing usually begins with a phenotype of interest such as a biological condition to derive a query signature, *i*.*e*., a set of differentially expressed genes. The query signature can be used to calculate similarities with compound perturbation data (also referred to as reference signatures) to indicate whether exposure to a specific compound is able to reverse or induce the phenotype of interest. In this process, a fundamental issue is to define the proper similarity between the query signature and the reference signature, which helps to elucidate the drug MOAs and identify new drug indications. Although various similarity measurements, including Cosine [6], KS [4], GSEA [5, 7], XSum [8], XCos [8], and sscMap [9], have been proposed to this end, a comprehensive study indicated that three issues remain. (1) All the existing similarity measurements are defined empirically in an unsupervised way. Due to the high dimensionality and the existence of high noise [10] in transcriptional signatures, it is difficult for empirically designed measurements to characterize the similarities between transcriptional signatures, resulting in inherently limited performance. (2) Most of the existing similarity measurements except for GSEA were developed specifically for Connectivity Map (CMap) [7]. Although the fashionability of CMap, its small scale restricts its application. Other resources, such as LINCS, which extended the CMap transcriptome data to a thousand-fold scale-up, have been proved to be much more useful. however, the data in these resources are heterogeneous. For example, the data in LINCS are profiled by the L1000 array platform, which is different from the microarray platform used by CMap [4, 5]. This inconsistency may make the existing methods fail to achieve generalized satisfactory performance. (3) The transcriptional signature of cells responding to a compound perturbation are affected by the cell type and the duration of treatment [11]. However, none of the existing methods consider this characteristic in a detailed and appropriate way for performing similarity calculations between transcriptional signatures, which may lead to incorrect analysis results.

In our study, we present Dr. Sim (**Sim**ilarity learning for **Dr**ug discovery), which is a general learning-based framework that automatically infers similarity measurement rather than being manually designed and can be used to characterize transcriptional phenotypic profiles for drug discovery with generalized good performance. For simplicity, we use “drug” and “compound” in our article to broadly refer to both FDA-approved chemicals, compounds and drug-like preclinical molecules. Basically, there are two main applications of such perturbation-based transcriptional profiles for phenotypic drug discovery: (1) drug annotation, *i*.*e*., elucidating MOAs for less well understood drugs; and (2) drug repositioning, *i*.*e*., proposing new indications for existing drugs [1-3]. Therefore, in our benchmark, we evaluated Dr. Sim on comprehensively publicly available *in vitro* and *in vivo* datasets in drug annotation and repositioning using high-throughput transcriptional perturbation data, and it was indicated that Dr. Sim significantly outperforms the existing strategies and is proved to be a conceptual improvement by learning transcriptional similarity to facilitate the broad utility of high-throughput transcriptional perturbation data for phenotypic drug discovery.

## Method

### General framework of Dr. Sim

Dr. Sim is a learning-based workflow designed to assign the label to a query transcriptional signature through calculating the similarity between the query transcriptional signature and each reference transcriptional signature cluster centroid using a measurement learned from reference transcriptional signatures rather than empirical design. The basic idea of Dr. Sim is a similarity learning schema that aims at making query signatures and reference signatures belonging to identical classes more similar, while making query signatures and reference signatures belonging to different classes more dissimilar [12]. In the benchmark of our current study, for illustration purposes, LINCS is used as the reference transcriptional signature data resource since it holds the largest-scale signatures produced by treating cancer cell lines with different compounds in various conditions [5]. Nevertheless, the application of Dr. Sim is not restricted to LINCS, and it can be applied directly to other transcriptional perturbation-based data resources for phenotypic drug discovery. Dr. Sim comprises three main steps: data preprocessing, model training, and similarity calculation.

### Data preprocessing

There are two steps in the data preprocessing stage: (1) quality control: Only transcriptional signatures treated by compounds for 6 hours (abbreviated as 6h) or 24 hours (abbreviated as 24h) in the nine cancer cell lines and two non-cancer cell lines in LINCS are retained because most experiments were performed under these conditions; (2) dataset splitting: The compound-induced expression signatures contain four attributes (cell type, compound, time point, and dosage) since they were defined as gene profile changes measured through comparing gene profiles collected after and before treating a cell line with a compound applied under a certain concentration, where the gene profile was calculated at a particular time point. We evaluate the influence of these four attributes on the transcriptional signatures (File S1). As demonstrated in Figures 1A-D, the cell type, compound, and time point substantially influence the distribution of transcriptional signatures, while the compound dosage does not. Therefore, to minimize the influence of the cell line and time point in calculating the similarities between compound-induced signatures, (i) the signatures in LINCS are split into 22 subsets according to cell type and time point (Figure 2A; eleven kinds of cell lines and two kinds of time points); (ii) the subset signature that has identical cell type and time point attributes to the query is used as a reference when a query search is conducted. Since the compound dosage slightly impacts the distribution of transcriptional signatures, the signatures treated with identical compounds at different dosages are considered replicates.

**Figure 1.** Visualization of the influence of the cell type, compound, time point, and compound dosage attributes on the expression signatures. In each subfigure, the data points differ in only one of the four attributes. **A**. Expression signatures profiled in the same cell line tend to cluster together, indicating that the cell type attribute impacts their distribution. **B**. Expression signatures profiled in the same compound tend to cluster together, indicating that the compound attribute impacts their distribution. **C**. Expression signatures profiled at the same time point tend to cluster together, indicating that the time point attribute impacts their distribution. **D**. Expression signatures profiled at different dosages tend to cluster together, indicating that the dosage attribute slightly impacts their distribution.

**Figure 2.** The Dr. Sim workflow. Dr. Sim comprises three main steps: data preprocessing, model training, and similarity calculation. **A**. In the first step, only signatures treated by compounds for 6h or 24h in the nine cancer cell lines and two non-cancer cell lines are retained, and the retained signatures are split into subsets according to the cell type and time point attributes. **B**. In the second step, Dr. Sim automatically infers a similarity measurement used for query assignment based on the training reference signatures. First, PCA is applied to the reference signatures to denoise and reduce dimensionality. A transformation matrix *P* is learned. Second, by applying LDA to the dimensionality-reduced signatures, a transformation matrix *L* is learned based on the signature labels indicating similarities and dissimilarities between signatures. The label of a signature is the compound class that it is induced by. Finally, the *TR* belonging to the identical class are median centered to derive the *TMR*. The *TR* is calculated using Eq. 1. C denotes the classes of signatures. **C**. In the third step, given a query signature, after transformation by *P* and *L*, its similarities to *TMR* are calculated by cosine similarity (Eq. 3). PCA, principal component analysis; LDA, linear discriminant analysis; *TR*, transformed references; *TMR*, transformed median-centered references.

### Model training

By adopting the linear discriminant analysis (LDA) metric learning algorithm (File S1) [12], Dr. Sim automatically infers a transcriptional similarity measurement used for query assignment based on the reference signatures (File S1). In the current study, the LDA algorithm was adopted since it achieves the best performance among all the metric learning algorithm (File S1) [12]. In summary, (1) principal component analysis (PCA) [13] is applied to the reference signatures to denoise and reduce dimensionality. A transformation matrix *P* is learned; (2) by applying LDA to the dimensionality-reduced signatures, a transformation matrix *L* is learned based on the signature labels indicating similarities and dissimilarities between signatures. The label of a signature is the compound class. The transformation matrix *L* that fits the relationships between signatures will project signatures into another space, in which signatures belonging to the identical class stay close to each other (intraclass similarity) while signatures belonging to different classes stay away from each other (interclass dissimilarity). In summary, the basic conception of LDA is to learn an optimal projection metric *L* that results in the optimal similarity measurement that aims at maximizing intraclass similarity and interclass dissimilarity; (3) the transformed references (*TR*) belonging to the identical class are median centered to obtain the transformed median-centered references (*TMR*; Figure 2C).

### Similarity calculation

For a query signature, after transformation by the *P* and *L* matrices, its similarities to the *TMR* are calculated by cosine similarity (File S1). The similarity score matrix is then ranked and can be used for query assignment. In drug annotation applications, the label of a query is assigned as the label of the reference that is most similar to the query since a positive score indicates compounds sharing a similar mechanism and activity [4, 5]. In drug repositioning applications, compounds that have the largest negative scores for the query are suggested since a negative score indicates that exposure to specific compounds can reverse the phenotype of interest [4, 5].

### Evaluating the rationale of Dr. Sim

Before evaluating the performance of Dr. Sim, we demonstrated the rationale of the designed workflow to make sure that it is appropriate for drug annotation and drug repositioning based on transcriptional data. First, we evaluate whether the similarity measurement learned automatically by Dr. Sim brings signatures with the same label close together while separating signatures with different labels. Second, we visualize the distribution of similarities between reference signatures and query signatures before and after applying Dr. Sim. For illustration purposes, the signatures having the most replicates in MCF7 at 24h are selected as an example. The selected signatures are split into queries and references at a ratio of 3:7, followed by applying Dr. Sim. As shown in Figures 3A-D, after transformation by the learned matrix *L*, signatures come from the same category have a tendency to cluster together, while signatures belonging to different classes tend to stay away from each other. To quantitatively measure this clustering tendency, we employed normalized mutual information (NMI) [14] from information theory in this study. If a clustering achieves a higher NMI, this indicates that it has higher intraclass similarity and lower interclass similarity. After transformation by the learned matrix *L*, the NMIs of the reference and query signatures were significantly higher in the 18 subset datasets (Figure 3E).

**Figure 3.** The evaluation of the rationale of Dr. Sim. **A**. Visualization of the clustering of reference signatures before transformation by the learned matrix *L*. **B**. Visualization of the clustering of reference signatures after transformation by the learned matrix *L*. Signatures belonging to the same class tend to cluster together, while signatures belonging to different classes tend to stay away from each other. **C**. Visualization of the clustering of query signatures before transformation by the learned matrix *L*. D. Visualization of the clustering of query signatures after transformation by the learned matrix *L*. Signatures belonging to the same class tend to cluster together, while signatures belonging to different classes tend to stay away from each other. **E**. After transformation by the learned matrix *L*, the NMIs of the reference and query signatures are significantly higher in the 18 subset datasets. **F**. Heatmap of the similarity between the reference and query signatures before transformation by the learned matrix *L*. **G**. Heatmap of the similarity between the reference and query signatures after transformation by the learned matrix *L*. The query signatures and reference signatures belonging to identical classes become more similar, while those belonging to different classes become more dissimilar. NMI, normalized mutual information.

Furthermore, we calculated the similarities between the reference signatures and query signatures. As shown in Figures 3F-G, query signatures and reference signatures belonging to identical classes become more similar, while query signatures and reference signatures belonging to different classes become more dissimilar. In other words, Dr. Sim maximizes the similarity between signatures if they share a similar expression pattern. The characteristics of Dr. Sim make it inherently suitable for drug annotation since in drug annotation, we assign a label to a query by searching for the most similar reference. Note that in drug repositioning, we intend to search for references that are most dissimilar to (maximally reverse) the query. To make the algorithm suitable for drug repositioning, we reversed the references before Dr. Sim trained a similarity measurement based on the references. In this case, Dr. Sim maximizes the similarity between signatures if they have opposite expression patterns.

## Results

### Evaluating the performance of Dr. Sim in drug annotation and drug repositioning

To demonstrate the advantage of Dr. Sim, we compared it against six other commonly used methods, including Cosine, KS, GSEA, XSum, XCos, and sscMap. NFFinder [15] and L1000CDS [16], which provide interactive websites to conduct similarity calculations, are not compared here because their core algorithms are KS and Cosine, respectively. The geneset size parameter, *i*.*e*., the total number of bottom- and top-ranked differentially expressed genes in the six methods, was set to 200, which is commonly used in other studies [17, 18]. The other parameters are set as default values. Note that Dr. Sim employs PCA to reduce data dimensionality. Hence, to prove that the performance improvement of Dr. Sim benefits from using LDA rather than PCA, we also compare Dr. Sim against a workflow that does not use LDA (referred to as no-LDA in our figures). In this study, Dr. Sim was evaluated in two scenarios, *i*.*e*., drug annotation and drug repositioning, based on comprehensively publicly available *in vitro* and *in vivo* transcriptional dataset.

### Scenario 1: Drug annotation

Compound perturbation-based transcriptional profiles have often been applied for drug annotation to uncover MOAs of less well understood drugs, such as how a drug produces an effect in the body. For example, the expression signature induced by thioridazine was found to have a strong similarity to those induced by DNA inhibitors, demonstrating that thioridazine exerts its anti-tumor activity by inhibiting DNA replication [19]. In the drug annotation benchmark scenario, the accuracy of predicting the MOAs of compounds was compared (File S1). The gold-standard MOAs of compounds in CMap and LINCS were retrieved from Huang et al. (Figure 4A; Table S1; File S1) [20]. As shown in Figure 4B, the accuracy of Dr. Sim is significantly higher and usually several times higher than that of the other methods. In addition, Dr. Sim outperformed the no-LDA workflow, indicating that the performance improvement of Dr. Sim benefits from employing the similarity learning-based strategy. In general, the training data size has an impact on the supervised learning model [21]. Therefore, we evaluated the impact of this factor on the accuracy of Dr. Sim. Not unexpectedly, as the training data size increases, the accuracy of Dr. Sim in predicting the MOAs of compounds increases (Figure 4C; File S1).

**Figure 4.** Benchmark of Dr. Sim against existing methods in drug annotation scenario. **A**. The proportion of compounds with MOA annotation information among all the compounds in CMap and LINCS. **B**. In predicting the MOAs of the compounds, Dr. Sim significantly surpasses all the existing methods in all the subset data. **C**. As the training data size increases, the accuracy of Dr. Sim increases, but that of the other methods does not. **D**. Dr. Sim surpasses all the existing methods based on heterogeneous transcriptional profiles. MOA, mechanisms of action; CMap, Connectivity Map; LINCS, Library of Integrated Network-Based Cellular Signatures.

Compound-induced profiles are usually heterogeneous. For example, expression profiles in LINCS are measured by L1000 technology due to cost constraints [5], while most of the expression profiles used by researchers to derive a query are measured by RNA-seq and microarray technology [22]. To demonstrate that Dr. Sim can achieve a generalized satisfactory performance in noisy and heterogeneous environments, heterogeneous signatures were used to predict the MOAs of compounds. Specifically, signatures in LINCS were used to predict the MOAs of compounds in CMap since the signatures in CMap and LINCS are profiled by microarray and L1000 technology, respectively, in a different way. As illustrated in Figure 4D, although compared with that in the homogeneous scenario, the accuracy of all the methods drops in the heterogeneous scenario, Dr. Sim still surpasses all the methods, demonstrating that its well-generalized performance is tolerant to noisy data and heterogeneity (Figures 4B-D).

### Scenario 2: Drug repositioning

The suggestion of novel indications for existing drugs, *i*.*e*., drug repositioning or repurposing, is significant for pharmaceutical study and is evolving as a method to reduce cost for drug discovery [23]. Compound-induced transcriptional profiles have been extensively applied in this area, reflecting a paradigm shift in pharmaceutical study, from the traditional seek of the magic bullet that targets a single pathogenic gene to the novel phenotypic means that inspect drug-gene-disease interactions from the system level. To comprehensively demonstrate the advantages of Dr. Sim in transcription-based drug repositioning, we benchmarked it with existing methods in three areas, *i*.*e*., *in vitro* and *in vivo* datasets and *in vivo* datasets with real-world evidence. We investigated whether Dr. Sim can suggest effective compounds for query signatures that are derived from the phenotype of interest in the three datasets. The validation workflow comprises three steps: collection of query signatures, compound scoring, and precision calculation (Figure 5A). More specifically, (1) we collected query signatures by comparing gene expression profiles from cancer cell lines or tumor tissues to the corresponding normal tissues; (2) by searching against reference signatures using the query signatures, the compounds were scored and ranked by Dr. Sim and the other methods. For each query signature, the *p*-value of a predicted compound was computed to classify the predicted compounds as effective or ineffective (File S1). (3) By collecting gold standard drug efficacy information from public resources, the precision metric, *i*.*e*., the proportion of real effective compounds among the predicted effective compounds, was calculated. Here, we focused on the precision metric rather than other metrics such as accuracy and recall because in real applications, it is practical to examine the few top indications; thus, we want the predicted effective compounds among the top suggestions to be truly effective as often as possible.

**Figure 5.** Benchmark of Dr. Sim against existing methods in drug repositioning scenarios. **A**. In drug repositioning, the validation workflow comprises three main steps: collection of query signatures, compound scoring, and precision calculation. (i) Query signatures were collected from CCLE, GTEx, and TCGA. (ii) Compounds were scored by Dr. Sim and the other methods. (iii) By collecting gold standard drug efficacy information from public resources, we calculated the precision metric. **B**. Dr. Sim surpasses all the existing methods in the *in vitro* dataset, indicating its high sensitivity. **C**. Dr. Sim achieves the highest nDCG score in BRCA and LUAD, demonstrating its superiority in the *in vivo* dataset. **D**. Of the patients predicted by Dr. Sim to respond to a drug, nearly all patients responded to it according to TCGA records. CCLE, Cancer Cell Line Encyclopedia; GTEx, Genotype-Tissue Expression; TCGA, the cancer genome atlas; nDCG, normalized discounted cumulative gain; BRCA, breast invasive carcinoma; LUAD, lung adenocarcinoma.

### Benchmarking Dr. Sim in the *in vitro* dataset for drug repositioning

For the *in vitro* dataset, the performance of predicting effective compounds against cancer cell lines was compared. We analyzed eight kinds of cancer cell lines in view of the availability of compound-induced signatures and compound efficacy information on these cancer cell lines (Table S2). Following the validation workflow,(1) query signatures were computed by comparing the expression profiles of cancer cell line in the Cancer Cell Line Encyclopedia (CCLE) [24] to the expression profiles of corresponding normal tissue in Genotype-Tissue Expression (GTEx; File S1) [25]; (2) compound efficacy data were collected from Genomics of Drug Sensitivity in Cancer (GDSC), ChEMBL, and Cancer Therapeutics Response Portal (CTRP), which are the largest publicly available compound efficacy databases (File S1) [26-28]. To minimize the time point attribute effect, the signatures profiled at 24h in LINCS are used as the reference signatures since drug efficacy in GDSC, ChEMBL, and CTRP is mostly measured at 24h or 72h. (3) For every query signature, compounds were scored and classified as effective or ineffective by each method (Figure 5A; File S1). As a result (Figure 5B), Dr. Sim obtained the highest precision. Most of the effective compounds predicted by Dr. Sim are truly effective.

### Benchmarking Dr. Sim in the *in vivo* dataset for drug repositioning

For the *in vivo* dataset, the performance in predicting FDA-approved drugs against diseases was compared. Three kinds of cancer were analyzed, *i*.*e*., lung adenocarcinoma (LUAD), breast invasive carcinoma (BRCA), and prostate adenocarcinoma (PRAD), based on the availability of FDA-approved drug reference signatures on these cancers (File S1). To further demonstrate the generalization of Dr. Sim, we apply it to a non-cancer disease, *i*.*e*., Alzheimer’s disease (AD). It should be noted the AD patient derived cell lines are not available in LINCS. Therefore, the signatures on the nine cancer cell lines in LINCS were used as references. Following the validation workflow, (1) query signatures were collected by comparing cancer and AD patient expression profiles to the corresponding normal tissue expression profiles (File S1); (2) FDA-approved drugs were downloaded from the National Cancer Institute (NIH) as the ground-truth drugs for performance validation. (3) For every query signature, drugs were scored and classified as effective or ineffective by each method. As depicted in Table S3, only Dr. Sim predicted FDA-approved drugs in BRCA, LUAD, and AD among all the tested methods, indicating its high sensitivity. To quantitatively compare the ranking result, the normalized discounted cumulative gain (nDCG) was adopted (File S1). Dr. Sim achieved the highest nDCG score in BRCA, LUAD, and AD (Figure 5C). Although none of the methods, including Dr. Sim, predicted FDA-approved drugs in PRAD, several top-ranked effective drugs predicted by Dr. Sim were validated *in vivo*. For example, calcitriol, the primary active metabolite of vitamin D, showed significant anti-neoplastic effect in preclinical models of PRAD (Table S3) [29].

### Benchmarking Dr. Sim in the *in vivo* dataset with real-world evidence for drug repositioning

For the *in vivo* dataset with real-world evidence, *i*.*e*., the clinical annotations, the performance in predicting drug response was compared. In view of the availability of drug response information in The Cancer Genome Atlas (TCGA) and the reference signatures on those drugs, BRCA and LUAD patients were analyzed (File S1). Following the validation workflow, (1) we collected 248 and 101 query signatures from BRCA and LUAD patients by comparing expression profiles from tumors to expression profiles from adjacent normal tissues. For BRCA patients, signatures at 6h and 24h in MCF7 were used as references. For LUAD patients, since two cell lineages of LUAD (A549 and HCC515) were profiled in LINCS, signatures in A549 and HCC515 were used as references. (2) The patients’ clinical outcomes were classified as “Response” or “Non-Response” (File S1); and (3) for each patient, the drugs were scored by each method. If the *p*-value of a drug was less than 0.01, we predicted that the patient would respond to the drug. As demonstrated in Figure 5D, Dr. Sim outperformed the other methods. Of the patients predicted by Dr. Sim to respond to a drug, nearly all the patients responded to it according to the TCGA records.

## Discussion

In the current study, we present a general learning-based framework, Dr. Sim, for transcriptional phenotypic drug discovery, in which the similarity measurement between signatures is learned from data rather than being manually designed. Traditionally, such similarity measurements have been defined in an unsupervised way, and due to the high dimensionality and the existence of high noise in these high-throughput data, they lack robustness with limited performance. For example, in previous benchmark studies, XCos and XSum performed best in drug annotation on the CMap dataset [8], while GSEA and sscMap performed best on the LINCS dataset [17]. The generalized robustness and superiority of Dr. Sim on different platforms and data sources were demonstrated on comprehensively *in vitro* and *in vivo* datasets. Taken together, Dr. Sim proves to be a conceptual improvement by learning transcriptional similarity to facilitate the broad utility of high-throughput transcriptional perturbation data for phenotypic drug discovery.

The cell response to a perturbation is affected by the cell type as well as the duration of treatment (Figures 1A-D). However, none of the existing methods consider this characteristic in a detailed and appropriate way for performing similarity calculations between transcriptional signatures, which may lead to an incorrect analysis result. It is shown that the accuracy of predicting the MOAs of compounds drops if we do not consider this characteristic (Figure S3). This may be explained by the fact that although perturbations that show similar activity across cell types exist, the activities of most perturbations are cell type and time point specific. These perturbations usually target specialized proteins. For example, glucocorticoid receptor agonists are maximum in cell lines in which the receptors of glucocorticoid are expressed [5]. In conclusion, cell type and time point attributes should be taken into consideration in calculating the similarity between transcriptional signatures.

The performance of Dr. Sim improves when the training data size increases (Figure 4C). At present, nearly one-third of the perturbation-based expression profiles in CMap and LINCS have few replicates due to cost constraints [4, 5]. With the decreasing cost of high-throughput sequencing, perturbation-based expression profiles are accumulating rapidly. It is conceivable that the performance of Dr. Sim can be further improved with such an increased amount of data.

Future improvements of Dr. Sim include (1) designing a more efficient similarity-learning algorithm to characterize transcriptional similarity [30]; and (2) identifying more efficient signatures through the genome-wide transcriptome, for example, the key pathway, to characterize the causality of diseases [31]. Then, such signatures could be incorporated into Dr. Sim for phenotypic drug discovery.

## Data availability

The processed MOAs of compounds in CMap and LINCS are available in the Table S1. The processed compound efficacy information is available in the Table S2.

## Code availability

Docker version of Dr. Sim can be installed at https://hub.docker.com/r/bm2lab/dr.sim/. The usage and manul of Dr. Sim is available at Github https://github.com/bm2-lab/DrSim/. The usage and manul of Dr. Sim is also available at BioCode at NGDC https://ngdc.cncb.ac.cn/biocode/tools/BT007273/.

## CRediT author statement

**Zhiting Wei**: Conceptualization, Data curation, Formal analysis, Investigation, Methodology, Writing – original draft, Validation, Writing – review & editing. **Sheng Zhu:** Data curation, Formal analysis, Methodology, Validation. **Xiaohan Chen:** Formal analysis, Visualization. **Chenyu Zhu**: Investigation, Visualization. **Bin Duan:** Investigation, Methodology. **Qi Liu**: Conceptualization, Writing – original draft, Validation, Writing – review & editing. All authors read and approved the final manuscript.

## Competing interests

The authors declare that they have no competing interest.

## Acknowledgements

This work was supported by the National Key R&D Program of China (Grant No. 2017YFC0908500, No. 2016YFC1303205), National Natural Science Foundation of China (Grant No. 31970638, 61572361), Shanghai Natural Science Foundation Program (Grant No. 17ZR1449400), Shanghai Artificial Intelligence Technology Standard Project (Grant No. 19DZ2200900) and Fundamental Research Funds for the Central Universities.

## Supplementary material

**File S1. Supplementary methods**

**Figure S1. Benchmarking the metric learning algorithm**

**A**. LDA is superior to NCA, LFDA, deepML and MLKR in terms of accuracy in drug annotation dataset. **B**. LDA is superior to NCA, LFDA, deepML and MLKR in terms of runtime efficiency in drug annotation dataset. LDA, linear discriminant analysis; NCA, neighborhood components analysis; LFDA, local fisher discriminant analysis; deepML, deep metric learning; MLKR, metric learning for kernel regression.

**Figure S2. The strategy of splitting training and testing data in drug annotation scenario**

For a kind of MOA, half of the compounds were used as reference compounds and half were taken as query compounds. The signatures induced by the reference compounds were used for training and the signatures induced by the query compounds were used for testing.

**Figure S3. The accuracy of predicting the MOAs of compounds when we do not take the impact of cell type and time-point attributes into consideration**

To exclude the impact of cell type, for example, signatures in MCF7 and A549 were used as query and references, respectively. To exclude the impact of time-point, for example, signatures at 6h and 24h in MCF7 were used as query and references, respectively.

**Table S1. The MOAs of compounds in CMap and LINCS**

**Table S2. The merged compound efficacy information for the eight cancer cell lines in LINCS**

**Table S3. (i-iii) The ranks of the FDA-approved drugs predicted by Dr. Sim and the other six methods in BRCA and LUAD; (iv) The ranks of drugs that have been validated in vivo in PRAD**

**Table S4. The FDA-approved drugs for BRCA, LUAD and PRAD**

## Notes

https://github.com/bm2-lab/DrSim

### Competing Interest Statement

The authors have declared no competing interest.

