## Supplementary File S1 for "Dr. Sim: Similarity Learning for Transcriptional Phenotypic Drug discovery"

**File S1. Supplementary information and methods**

**General framework of Dr. Sim**

**Data preprocessing**

LINCS L1000 level-4 transcriptional signatures are calculated by comparing compound-induced profiles to control. All the associated metadata were downloaded from National Center for Biotechnology Information Sequence Read Archive (GSE92742) and LINCS data portal 2.0 [1]. The raw signatures stored in GCTx format were converted to human-readable format by the cmapPy package [2]. To visualize the influence of the cell type, compound, time point and dosage attributes on the distribution of compound-induced transcriptional signatures, we downloaded the processed transcriptional signatures from the L1000FWD website [3]. L1000FWD reduces the dimensionality of raw transcriptional signatures using principal component analysis (PCA), followed by performing t-distributed stochastic neighbor embedding (tSNE) on them. For illustration purposes, downloaded processed signatures having most replicates are selected as an example. As demonstrated in Figures 1A-D, the cell type, compound and time point greatly impact the distribution of signatures, while the compound dosage does not.

**Model training and similarity calculation**

Dr. Sim formulates a metric learning-based framework in which the similarity measurement used for query assignment is learned from the reference data rather than being manually designed like Cosine, KS, GSEA, XSum, XCos and sscMap. Briefly, PCA was first applied to the reference data to denoise and reduce dimensionality. A transformation matrix *P* is learned. Then, by applying LDA [4] to the dimensionality-reduced signatures, a transformation matrix *L* is learned based on the signature labels indicating similarities and dissimilarities between signatures. The compound class is used as the training label fed to LDA. In summary, the basic idea of LDA is to learn an optimal transformation matrix *L* that leads to the optimal similarity measurement that aims at maximizing intraclass similarity and interclass dissimilarity. The transformed references denoted as *TR* can be calculated as follows:

$TR=R*P*L$	(1)

Where *R* is the reference signatures. The *TR* belonging to the identical class are median centered to obtain the transformed median-centered references (denoted as *TMR*)*.* For a query signature Q, we transformed it as follows:

$TQ=Q*P*L$	(2)

Where *P* and *L* are learned by PCA and LDA based on the reference signatures. Lastly, the similarities between the query signature and the *TMR* are computed by cosine similarity as follows:

$similarity=\frac{TMR*TQ}{\left| \left| TMR \right| \right| \left| \left| TQ \right| \right|}$	(3)

The n_components parameter in the PCA algorithm implemented in scikit-learn [5] was set to 0.98 (select the number of components so that the amount of explained variance in the reference signatures is greater than 98 percentage). The n_components parameter in the LDA algorithm implemented in scikit-learn [5] was set to 50. Other parameters were set to default values.

**Benchmarking the metric learning algorithm**

To select the most appropriate metric learning algorithm, the performance of the commonly used metric learning algorithms, including linear discriminant analysis (LDA), neighborhood components analysis (NCA), local fisher discriminant analysis (LFDA), deep metric learning (deepML, <https://github.com/KevinMusgrave/pytorch-metric-learning>) and metric learning for kernel regression (MLKR) were compared in drug annotation dataset [6]. The drug annotation dataset rather the drug repositioning dataset was used to benchmark their performances since drug annotation gold standard was well defined (see ***methods***). The large margin nearest neighbor (LMNN) algorithm was not considered since it takes too much time. The *n_components* parameter in LDA, NCA, LFDA and MLKR is set to 50. Other parameters are set to default values. In deepML, the loss function and the optimizer are set to *TripletMarginMiner* and *Adam,* respectively. Other parameters are set to default values. As demonstrated in Figures S1A-B, LDA is superior to other methods in terms of accuracy and runtime efficiency. Therefore, LDA was adopted as the final metric learning algorithm in the current study.

**Calculation of the NMI**

In [probability theory](https://en.wikipedia.org/wiki/Probability_theory) and [information theory](https://en.wikipedia.org/wiki/Information_theory), normalized mutual information (NMI) [7] is used for measuring amount of information of a [clustering](https://en.wikipedia.org/wiki/Cluster_Analysis). To calculate the NMI of a collection of signatures, a k-means clustering model was constructed to predict their clustering labels. The NMI was calculated by comparing the predicted clustering labels with the true clustering labels of the given signatures. The *k* parameter in the k-means clustering model is set to the number of classes of the given signatures. Other parameters were set to default values.

**Test scenario 1: Drug annotation**

**Collection of compound MOA information**

The MOAs of compounds in CMap and LINCS were retrieved from Huang et al. [8]. The authors manually curated MOA tag descriptions for all compounds. In total, there are 213 kinds of MOA tag descriptions (Table S1). Among the 24036 compounds in CMap and LINCS, only 2597 compounds have explicit MOA annotation.

**Calculation of the accuracy of predicting the MOAs of compounds**

The accuracy of predicting the MOAs of compounds was compared on the 22 subset data. For every subset data: (1) Compounds inducing not less than five signatures and have explicit MOA annotation were kept. To make sure that compounds in the queries are new to references, the following strategy was adopted in splitting subset data into queries and references (Figure S2). For a kind of MOA, half of the compounds were used as reference compounds and half as query compounds. The signatures induced by the reference compounds were used for training and the signatures induced by the query compounds were used for testing. (2) For a query, its similarities to the references were calculated using Dr. Sim, the no-LDA workflow and the six other methods. The MOA of a query is then assigned as the MOA of the reference compounds that is most similar to the query. (3) At last, we calculate the accuracy, *i.e.*, the proportion of correctly predicted queries among all the queries. To avoid uncertainty in splitting reference and query compounds, this procedure was repeated ten times and the average accuracy was used as the final result.

**Evaluation of the influence of training data size on Dr. Sim**

To evaluate the impact of the training data size on the performance of Dr. Sim and the six other methods, the following experiments were performed. We selected compounds inducing not less than 10 signatures in each subset data. The signatures were split into queries and references as mentioned above. In the high replication scenario, all the signatures inducing by a compound were used for training. In the low replication scenario, only half of the signatures inducing by a compound were used for training. Finally, we compared the performances of Dr. Sim between the high and low replication scenarios.

**Collection of query signatures from CMap**

CMap produces genome-wide transcriptional profiles from human cell lines treated with compounds at different dosages mainly for 6H [9]. The current version (build 02) of CMap contains 7056 transcriptional profiles, including 6100 induced by treating cell lines with 1309 compounds and 956 induced by treating cell lines with DMSO (control). CMap raw Affymetrix expression data and their annotations were downloaded from <https://portals.broadinstitute.org/cmap>. Only transcriptional profiles measured in MCF7 and PC3 (among the LINCS eleven cell lines) at 6H (among the LINCS two time points) were kept. Expression data were fitted and normalized using the affyPLM [10] and affy [11] Bioconductor packages. The Probe IDs in microarray were converted to gene IDs using the hthgu133a Bioconductor package. Expression levels of multiple probes matching to the same gene were averaged as the expression level of that gene. Only genes measured both in CMap and LINCS platforms were retained. Finally, following the pipeline in LINCS, the robust z-scoring metric [12] was employed to derive query signatures as follows: the differential expression of gene *x* in the *i* sample within a batch were computed as follows:

$z_{i}= \frac{x_{i}-median\left( X \right)}{1.4826*MAD \left( X \right)}$	(4)

Where *X* is the vector of normalized gene expression of gene *x* across all control samples within that batch, *MAD* is the median absolute deviation function, and the factor of 1.4826 makes the denominator a consistent estimator of scale for normally distributed data.

**Test scenario 2: Drug repositioning**

**Calculation of the *p*-value of a compound**

To classified a predicted compound as effective or ineffective against a query, the *p*-value of the compound was computed by borrowing the idea from Subramanian et al. [12]. Briefly, the similarity score *s* between the compound and the query is calculated by Eq. 3. Then, the similarity score *s* is compared with the background similarity score *S* that is calculated between the compound and a compendium of random queries. The random queries are generated by randomly shuffled the query 1000 times. The background similarity score *S* is calculated by Eq. 3. Finally, the *p*-value of the compound is calculated as follows:

$p= \frac{\sum_{i=1}^{1000} S_{i}\leq s}{1000}$	(5)

**Collection of the in-vitro data**

The Cancer Cell Line Encyclopedia (CCLE) [13] contains gene transcriptional profiles from more than 1000 human cancer cell lines. We analyzed nine cancer cell lines among them, based on the availability of compound-induced signatures on these cancer cell lines in LINCS. The nine cancer cell line transcriptional profiles were downloaded from <https://portals.broadinstitute.org/ccle/>. Their corresponding normal tissue transcriptional profiles were downloaded from Genotype-Tissue Expression (GTEx) [14], which collected transcriptional profiles from 54 non-diseased tissue sites across nearly 1000 individuals. For every cancer cell line, its transcriptional profile was merged with its corresponding normal tissue transcriptional profiles. The merged transcriptional profiles were then normalized using edgeR [15] Bioconductor packages. Genes not detected in LINCS L1000 technology were filtered. Following the pipeline in LINCS, the robust z-scoring metric [12] was employed to derive the query signature for each cencer cell line (calculated using Eq. 4).

To be comprehensively, the gold standard drug efficacy information was downloaded from Genomics of Drug Sensitivity in Cancer (GDSC) [16], ChEMBL [17], and Cancer Therapeutics Response Portal (CTRP) [18]. GDSC and ChEMBL quantitatively measure drug efficacy against cancer cell lines using half-maximal inhibitory concentration (IC50) metric while CTRP measures drug efficacy using the area under concentration-response (AUC) metric. In GDSC and ChEMBL, if the IC50 of a compound in a cell line is greater than 10000nM, the compound is considered ineffective against the cell line, otherwise is considered effective [19]. In CTRP, if the AUC of a compound in a cell line is greater than the median AUC of all compounds in that cell line, the compound is considered as ineffective, otherwise is considered as effective [18]. For a compound having multiple IC50 or AUC in the three databases, we used the median to summarize them. Finally, compound efficacy information in GDSC, ChEMBL, and CTRP was merged. Compounds between LINCS and the three databases were mapped using the compound generic name. The compound efficacy information is available in the supplementary file (Table S2).

**Collection of the in-vivo data**

In the in-vivo validation scenario, the FDA-approved drug information downloaded from National Cancer Institute (NIH) was used as the ground-truth for performance evaluation. Downloaded drugs were mapped to LINCS using the drug generic name. We collected RNA-seq transcriptional profiles of four kinds of cancers (including BRCA, LUAD, PRAD and SKCM) and their corresponding adjacent normal tissues from TCGA since only those cancers have FDA-approved drug-induced reference signatures in LINCS. SKCM was not for further analysis since there is not enough normal tissue transcriptional profile for the calculation of the query signature. The AD patient processed transcriptional profiles were downloaded from NCBI GEO (GSE26972). Samples from 3 female non-demented controls and 3 female AD patients were included in this study. Expression levels of multiple probes matching to the same gene were averaged as the expression level of that gene. Only genes measured in LINCS platforms were retained. Following the pipeline in LINCS, for every cancer type and AD, the robust z-scoring metric was employed to derived query signature (calculated using Eq. 3). The gene Ensemble ID was converted to Entrez ID using clusterProfiler Bioconductor packages [20]. Genes not detected in the LINCS L1000 platform were filtered. Two cell lineages of LUAD (A549 and HCC515) and PRAD (PC3 and VCAP) are profiled in LINCS. Therefore, for LUAD, signatures at 6H and 24H in A549 and HCC515 were used as reference signatures; for PRAD, signatures at 6H and 24H in PC3 and VCAP were used as reference signatures; for BRCA, signatures at 6H and 24H in MCF7 were used as reference signatures. For AD, the signatures on the nine cancer cell lines in LINCS were used as references since the AD patient derived cell lines are not available. The predicted results at 6H and 24H were merged directly and sorted by the similarity score Table S3. The FDA-approved drugs information used in this article is available in the Table S4.

**Collection of the in-vivo data with real-world evidence**

A part of patients in TCGA received drug treatment. The drug response records were used as the ground-truth to evaluate the performance of predicting drug response. The “Complete response” and “Partial response” in the records were classified as “Response” and the “Clinical progressive disease” and “Stable disease” in the records were classified as “Non-response”. BRCA and LUAD patients were analyzed since only drugs that treat BRCA and LUAD patients have reference signatures in LINCS. In total, 248 and 101 query signatures from BRCA and LUAD patients were available by comparing transcriptional profiles from tumors to those from adjacent normal tissues. For BRCA patients, signatures at 6H and 24H in MCF7 were used as references. For LUAD patients, since two cell lineages of LUAD (A549 and HCC515) were profiled in LINCS, signatures at 6H in A549 and HCC515 were used as references (signatures were not available at 24H in A549 and HCC515 for those drugs in TCGA records).

**The normalized discounted cumulative gain**

Normalized discounted cumulative gain (nDCG) is a measure of the ranking quality. In the current study, nDCG measures the gain of an FDA-approved drug based on its rank position in the result list. The gain is accumulated from the top of the result list to the bottom, with the gain of each result discounted at lower ranks. nDCG is calculated as follows:

$nDCG= \frac{\sum_{i=1}^{j} \frac{1}{{log}_{2}(R(i)+1)}}{\sum_{i=1}^{N} \frac{1}{{log}_{2}(R(i)+1)}}$	(6)

Where *j* is the number of FDA-approved drugs predicted by Dr. Sim and the other commonly used methods, *N* is the number of predicted effective drugs, *R(i)* is the rank of a predicted drug.
