## Supplementary figures and images for "Dr. Sim: Similarity Learning for Transcriptional Phenotypic Drug discovery"

### Figure 1

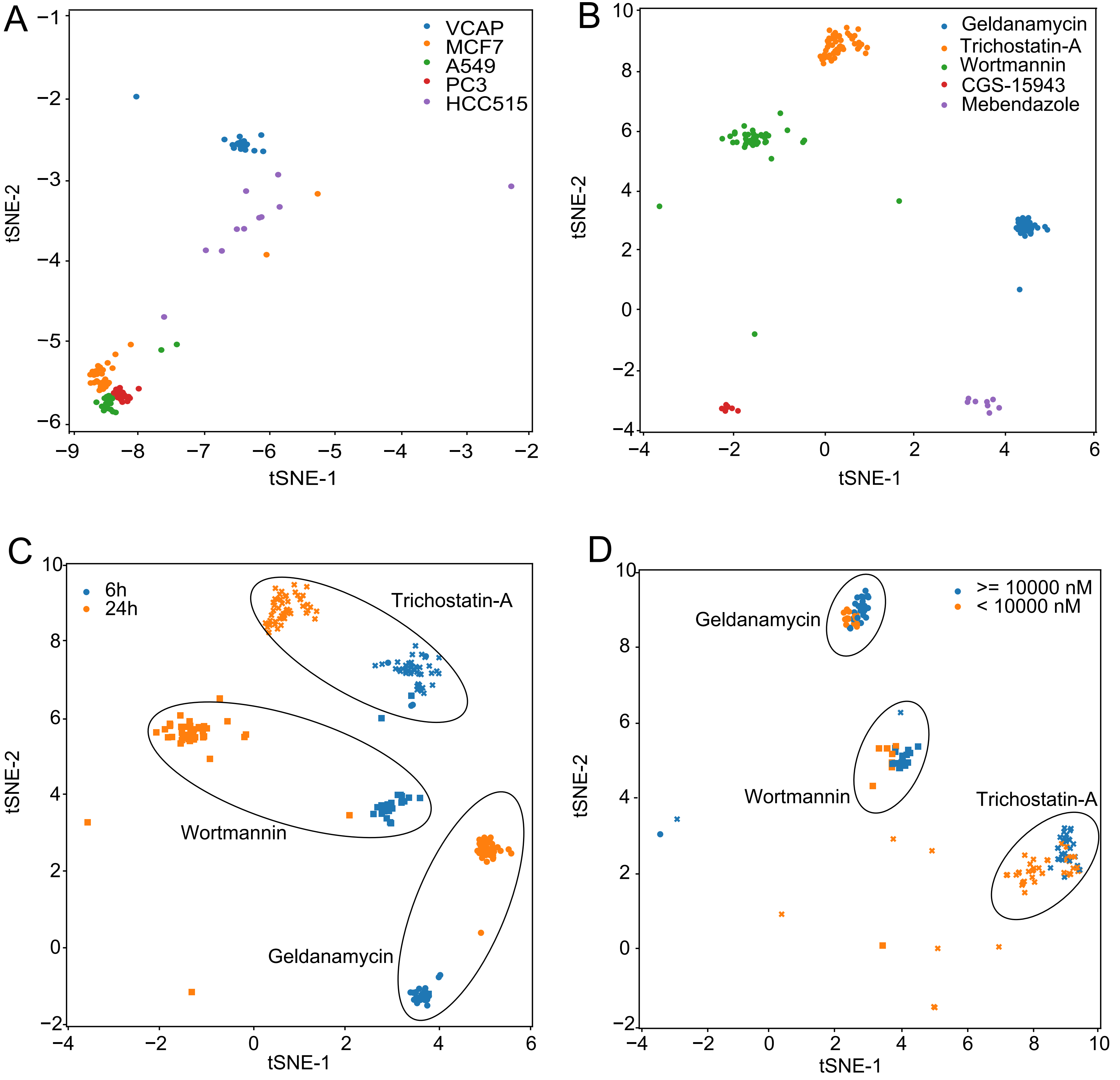

### Figure 2

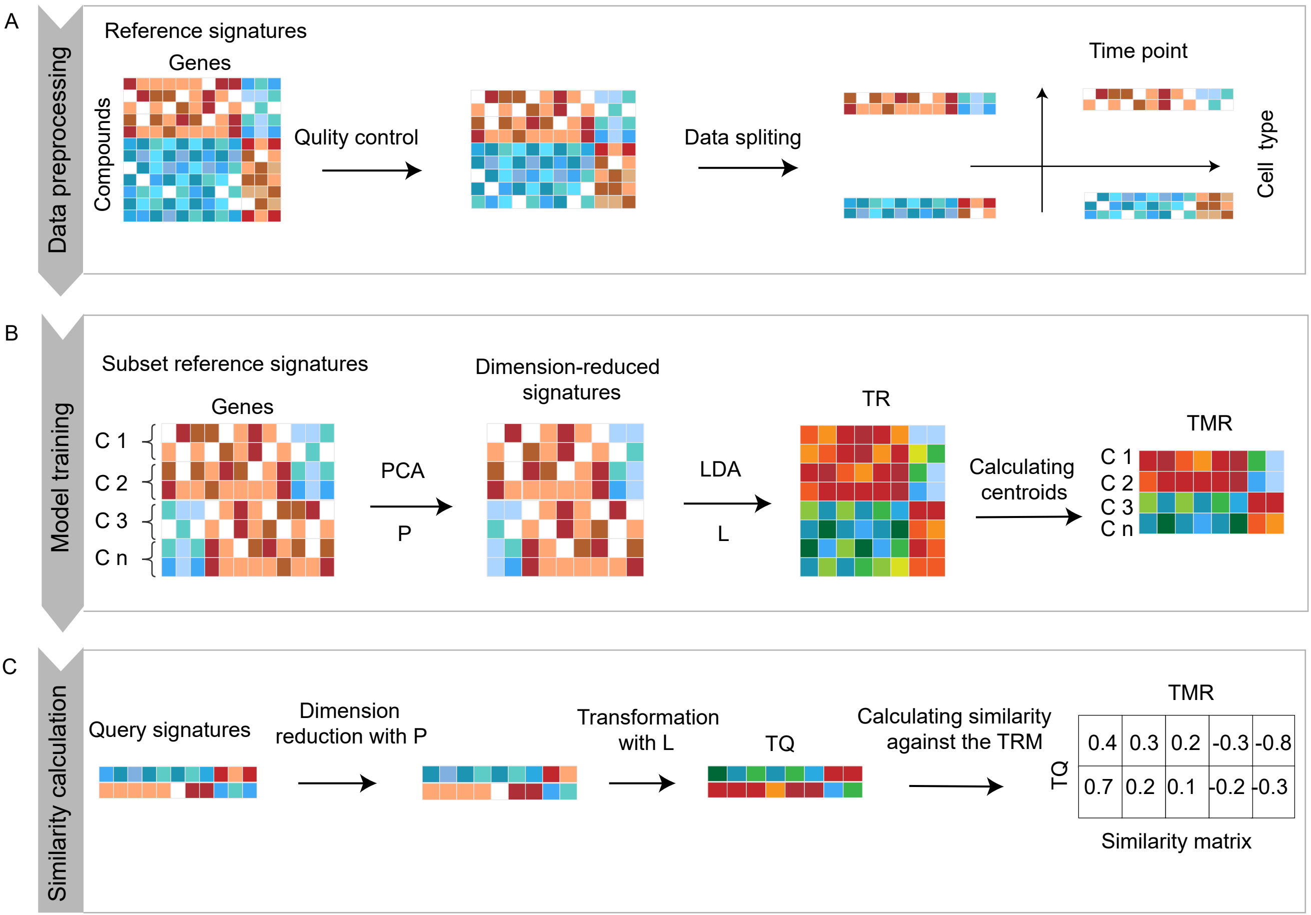

### Figure 3

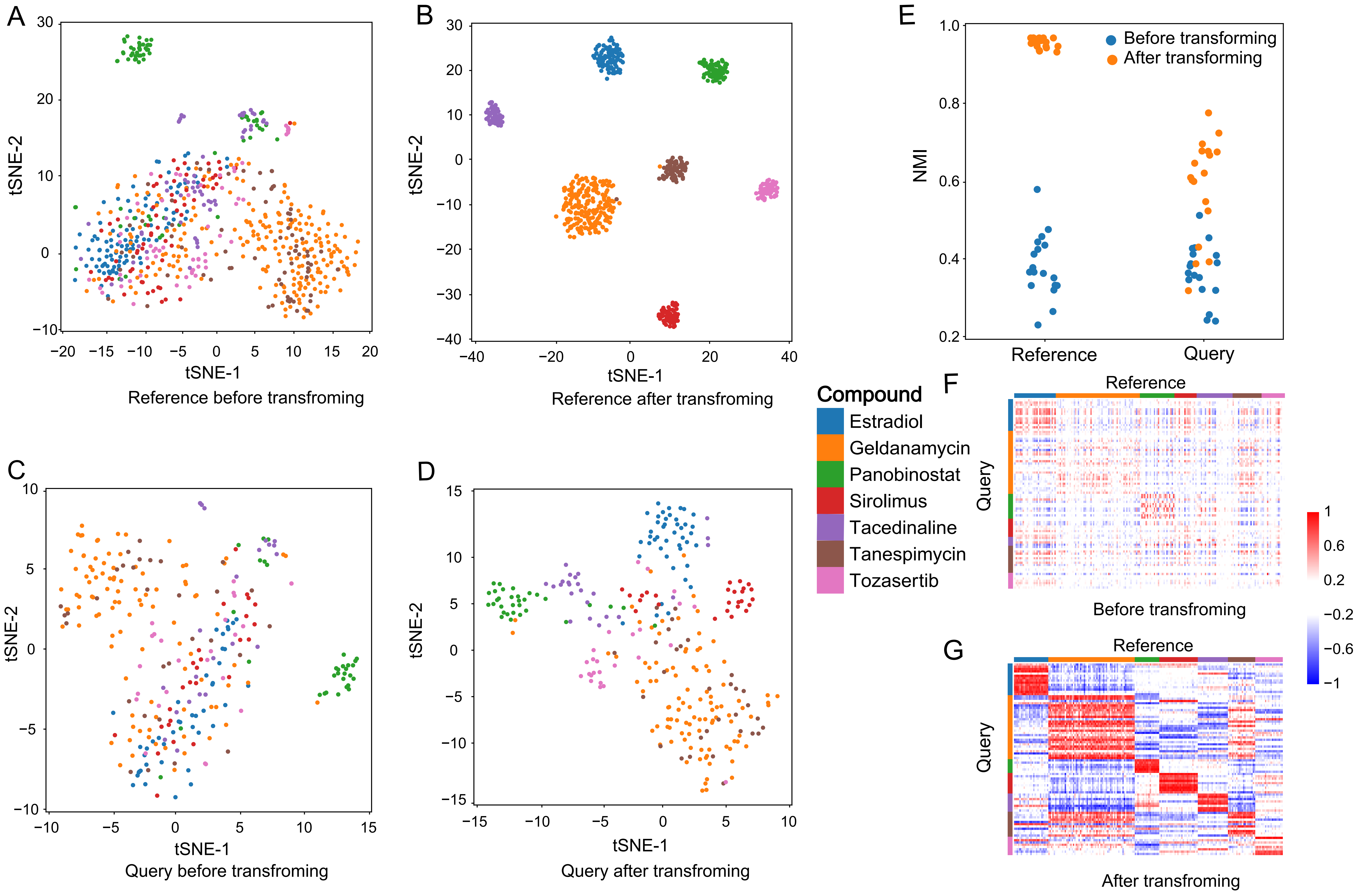

### Figure 4

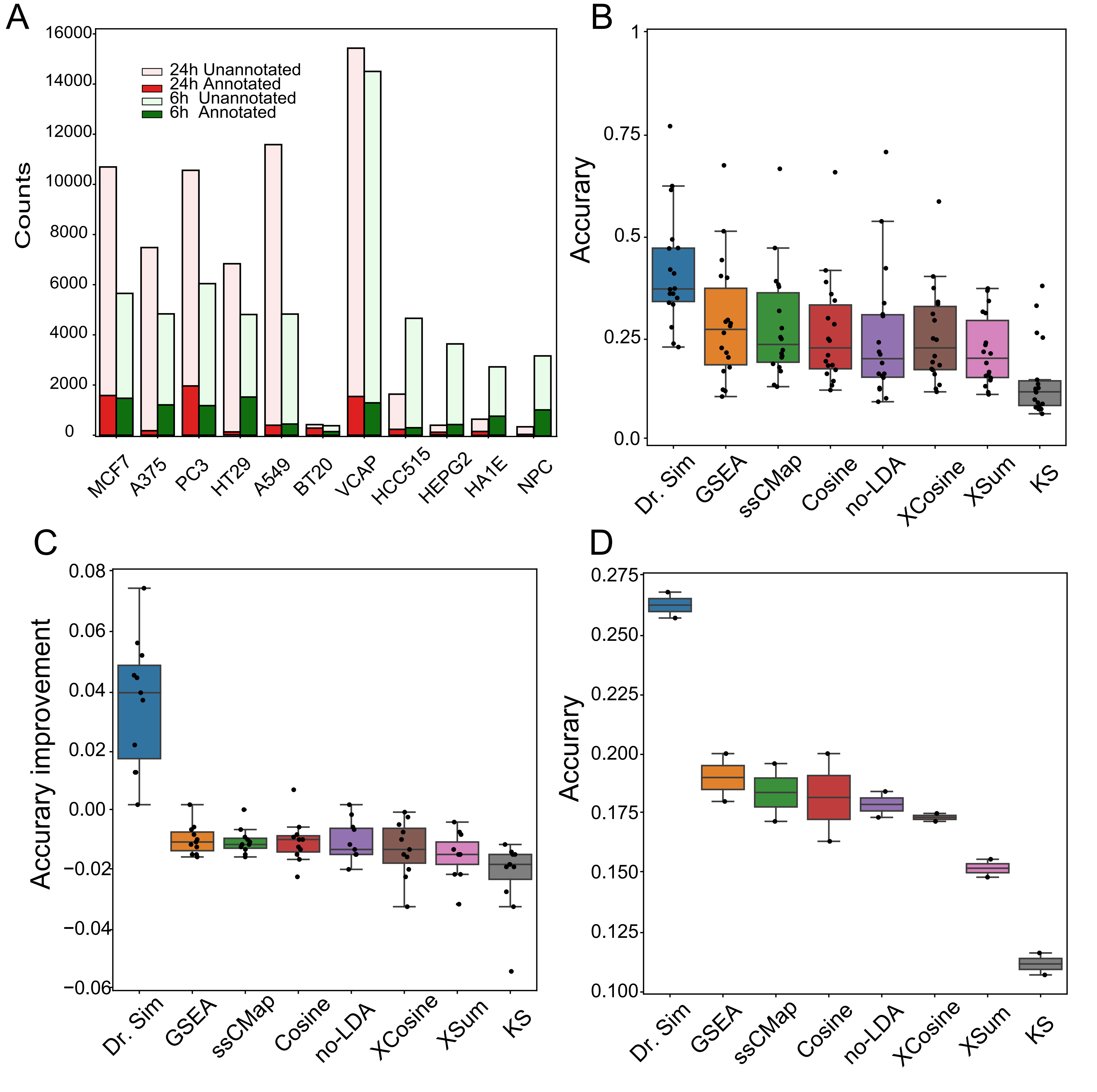

### Figure 5

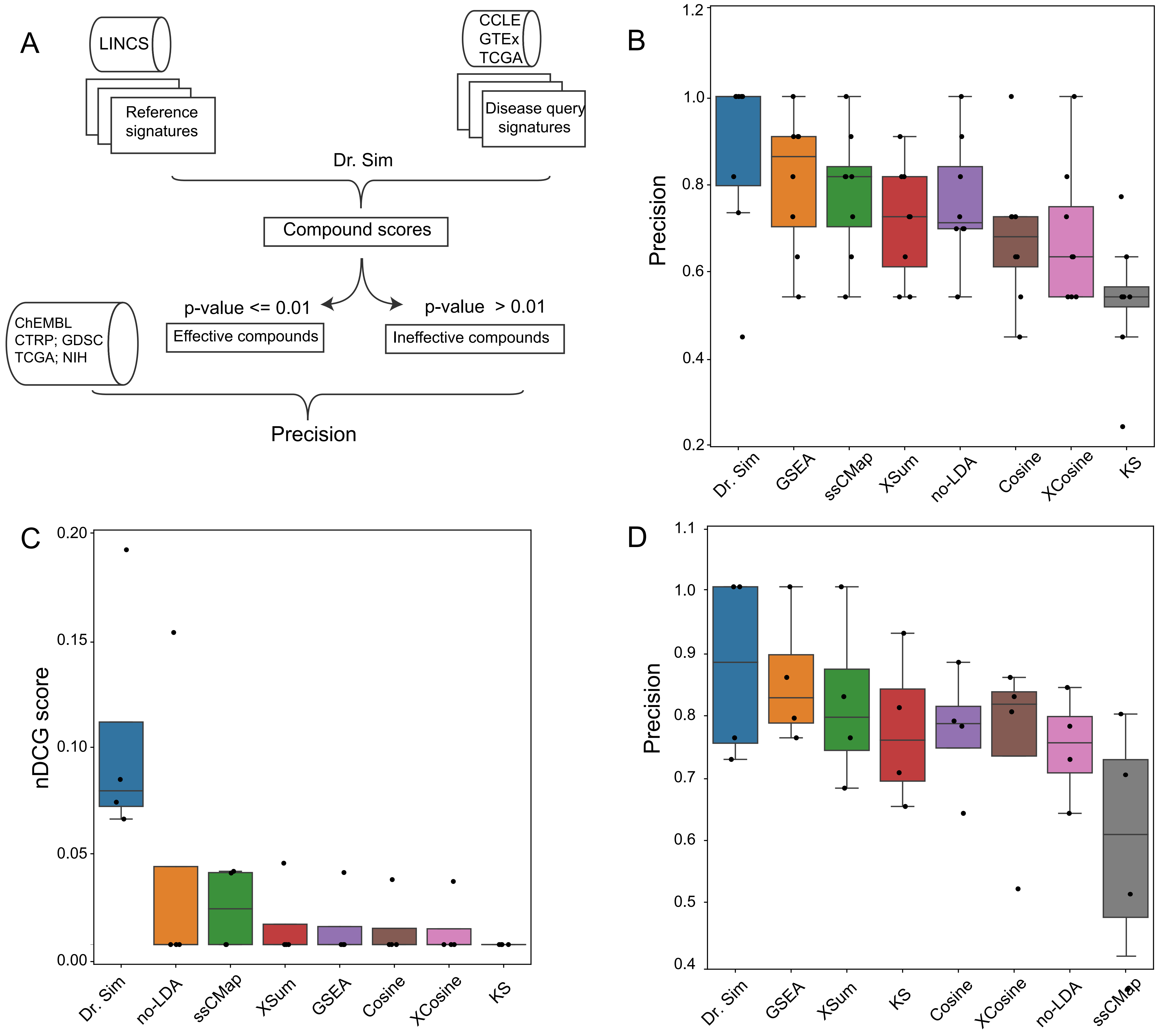

### Supplementary Figure S1

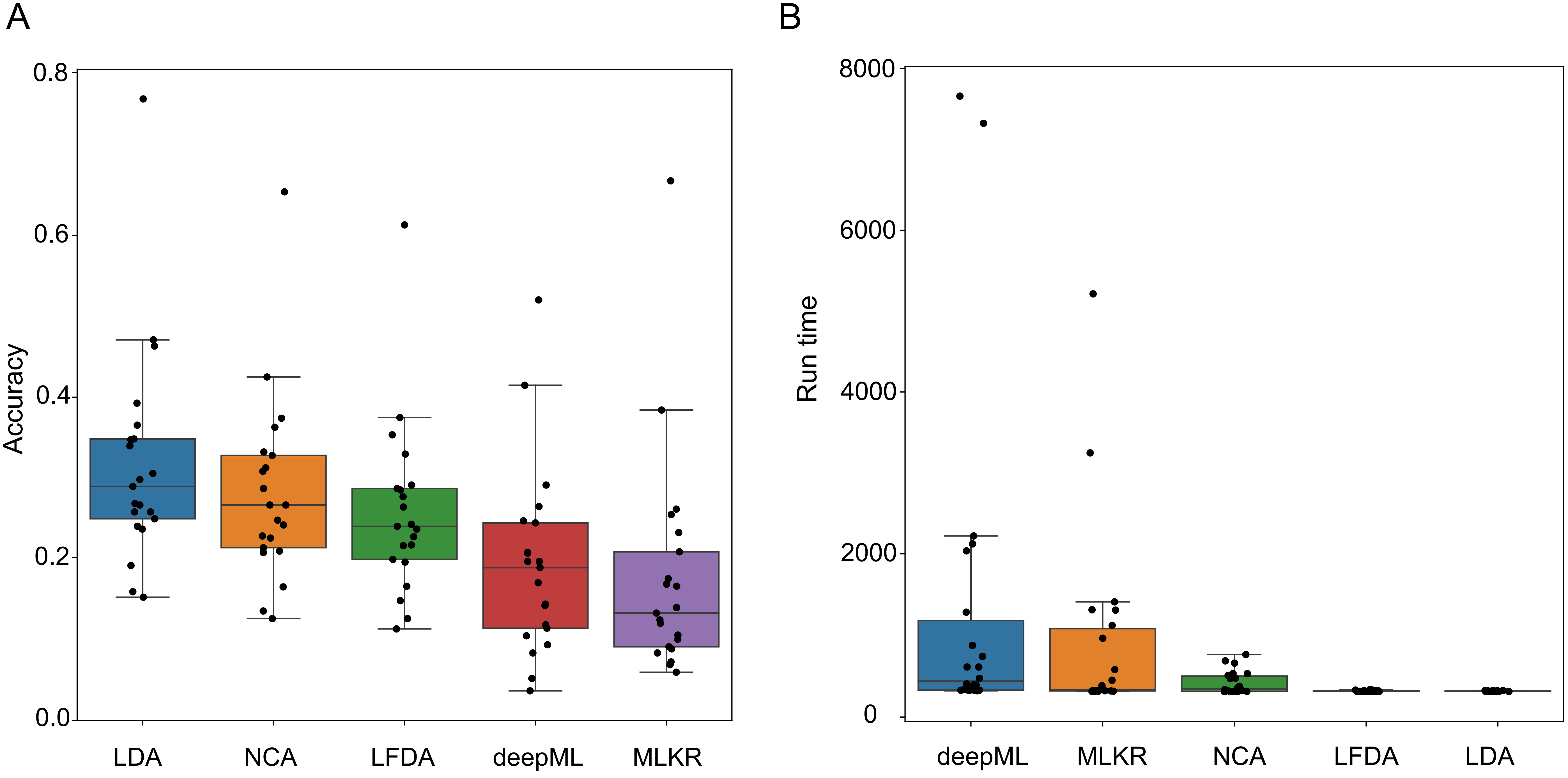

### Supplementary Figure S2

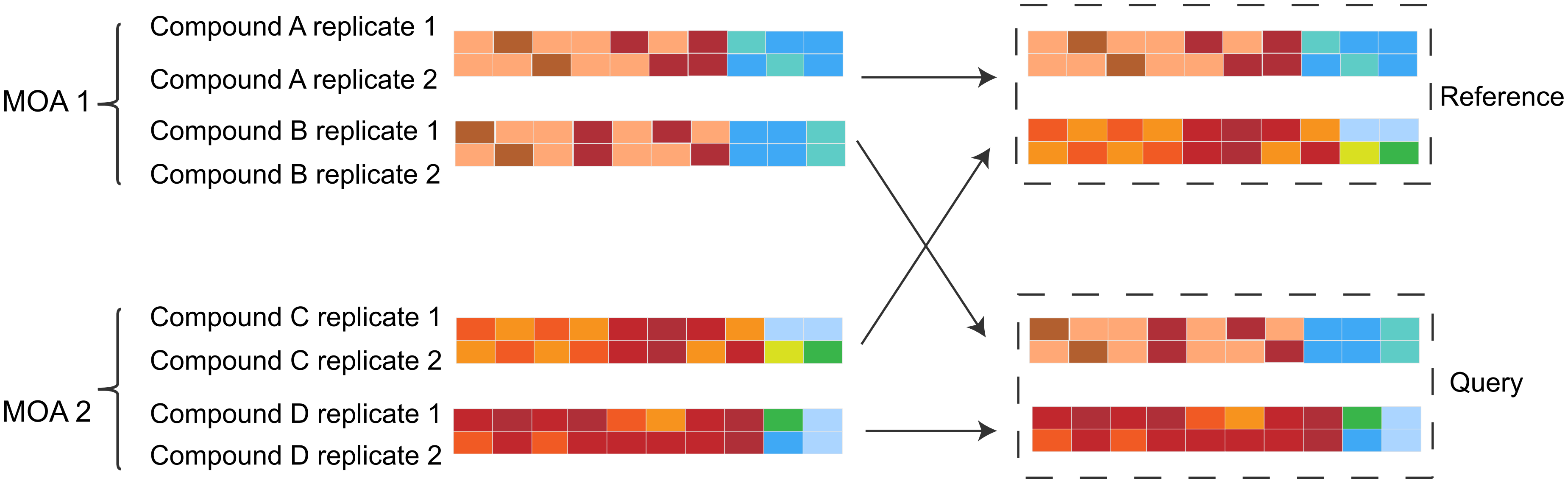

### Supplementary Figure S3

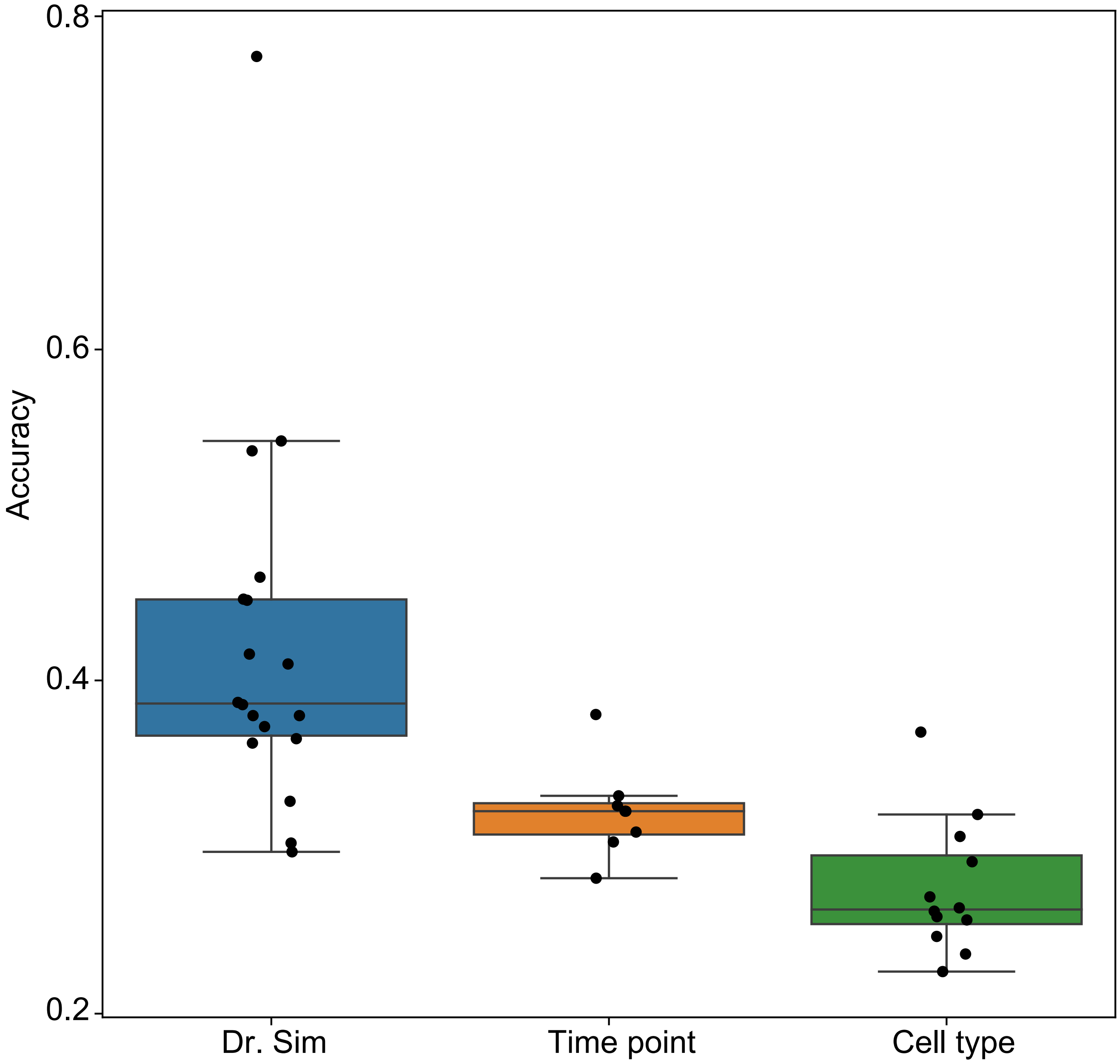
